# Functional analysis of the chromosomal passenger complex in Arabidopsis

**DOI:** 10.1101/2020.03.19.998880

**Authors:** Shinichiro Komaki, Hidenori Takeuchi, Yuki Hamamura, Maren Heese, Takashi Hashimoto, Arp Schnittger

## Abstract

A key regulator of cell division in all eukaryotes is the kinase Aurora B, which is encoded by the Aurora 3 (AUR3) gene in Arabidopsis. Aurora B has at least two central functions during cell division. On the one hand, it is essential for the correct, i.e. balanced, segregation of chromosomes in mitosis and meiosis by controlling kinetochore function. On the other hand, Aurora B acts at the division plane, where it is necessary to complete cytokinesis. To accomplish these two spatially distinct functions, Aurora B in animals is guided to its sites of action by Borealin, INCENP, and Survivin that build together with Aurora B the chromosome passenger complex (CPC). However, besides Aurora homologs, only a candidate gene with restricted homology to INCENP has so far been described in Arabidopsis raising the question whether there exists a full complement of the CPC in plants and how Aurora homologs are targeted subcellularly. Here, we have identified and functionally characterized a Borealin homolog, BOREALIN RELATED (BORR), in Arabidopsis. This, together with detailed localization studies including the putative Arabidopsis INCENP homolog, supports the existence of a CPC in plants.

## INTRODUCTION

Equal chromosome segregation during cell division is crucial for survival, growth, and reproduction of every organism. Chromosome segregation is assured by the M-phase checkpoint, which involves two regulatory units, the spindle assembly checkpoint (SAC) and the chromosomal passenger complex (CPC) (Carmena et al., 2012).

Chromatids are held together by Cohesin, a proteinaceous ring-like structure, which needs to be cleaved by Separase to allow their separation in anaphase. The SAC inhibits the Anaphase Promoting Complex/Cyclosome (APC/C), an E3 ubiquitin ligase, which mediates the degradation of Securin, an inhibitor of Separase, until all kinetochores are properly attached to microtubules of opposing poles (London and Biggins, 2014; Komaki and Schnittger, 2016).

The CPC has been well characterized in yeast and mammals, where it has been found to consist of four proteins: Aurora kinase B (Aurora B), Borealin, Inner centromere protein (INCENP), and Survivin. The CPC fulfills several functions during mitosis. In particular, it is involved in ensuring that all kinetochores are attached to microtubules emanating from opposing poles (Kitagawa and Lee, 2015). The CPC localizes to inner centromere where it activates Aurora B in response to low inter-kinetochore tension. Active Aurora B phosphorylates kinetochore proteins leading to the destabilization of erroneous microtubule attachments. Once proper kinetochore-microtubule attachments are established from opposing poles, which gives high inter-kinetochore tension, Aurora B is spatially separated from kinetochores resulting in proper bistable spindle formation.

Aurora B belongs to the Aurora kinase family of serine/threonine kinases that are highly conserved in the eukaryotic kingdom (van der Waal et al., 2012; Weimer et al., 2016). While yeast possesses a single Aurora homolog, mammals have three Aurora kinases, Aurora A, Aurora B, and Aurora C, among which only Aurora C acts in meiosis (Goldenson and Crispino, 2015). Since these three kinases share a common consensus phosphorylation motif, it is thought that interacting proteins are important for their localization and substrate specificity.

Aurora A interacts with the spindle assembly factor TPX2, and localizes to spindle microtubules to regulate spindle assembly (Gruss and Vernos, 2004). In contrast, Aurora B and Aurora C, the catalytic subunit of the CPC, is involved in the correction of erroneous kinetochore-microtubule attachments, activation of the SAC, and cytokinesis in mitosis and meiosis. These diverse functions are based on its dynamic localization pattern controlled by the three non-catalytic subunits of the CPC (van der Horst and Lens, 2014).

INCENP is the largest non-catalytic subunit of the CPC, which in animals and yeast directly binds to all other components of the complex. Borealin and Survivin interact with the conserved N-terminal region of INCENP while Aurora B binds to the C-terminal domain of INCENP, called IN-box (Carmena et al., 2012). The IN-box is required for interaction with and activation of Aurora B (Honda et al., 2003).

While the N-terminus of Borealin acts as the INCENP-binding region its C-terminus contains a homo-dimerization domain that is involved in a stable CPC localization at centromeres (Bekier et al., 2015). In addition, the phosphorylation status of the central part affects the centromere localization and steady-state level of Borealin itself (Kaur et al., 2010; Date et al., 2012).

Aurora kinases in plants are categorized into two groups, alpha-Aurora and beta-Aurora, based on the localization pattern and sequences (Weimer et al., 2016). Interestingly, these two groups have mixed features of the animal Aurora A and Aurora B/C groups. For instance, AUR1 and AUR2 in Arabidopsis, both belonging to the alpha class, localize to spindle microtubules, which is reminiscent of Aurora A, but they also localize to the central region of phragmoplasts paralleling Aurora B/C at the cell cleavage site. AUR3, a member of the beta-Aurora group, localizes to kinetochores, similar to Aurora B (Komaki and Schnittger, 2017), but in contrary to Aurora B, it does not accumulate at the division plane (Demidov et al., 2005).

While *aur1* and *aur2* single mutants do not show any obvious growth alterations, the *aur1 aur2* double null mutant is gametophytic lethal (Van Damme et al., 2011). A weak loss-of-function *aur1 aur2* double mutant exhibits altered division plane orientation, reduced pollen viability, and enhanced vascular cell differentiation. These defects can be rescued by expressing either AUR1 or AUR2 but not AUR3, indicating that alpha-Auroras and beta-Auroras have distinct functions (Van Damme et al., 2011; Demidov et al., 2014; Lee et al., 2019).

Given the sparse information about the non-catalytic subunits of the plant CPC, it is not even clear whether there is a conserved CPC function in plants. The putative homolog of INCENP in Arabidopsis has a long N-terminal region of unknown function, which is conserved only in plants. Although Arabidopsis *incenp* mutants, also known as *wyrd* (*wyr*), show defects in gametophytic cell division (Kirioukhova et al., 2011), it is still unclear whether Arabidopsis INCENP acts as part of a putative plant CPC because of missing information about its localization and binding partners.

Here, we present the identification and functional characterization of an Arabidopsis Borealin homolog, which colocalizes with the INCENP homolog to the inner centromere and the central domain of the phragmoplast. We also observed that only AUR3 acts as the catalytic subunit of the plant CPC. These data underscore the mixed features of plant Aurora kinases.

## RESULTS

### Identification of a putative Borealin homolog in plants

As a first step to determine whether plants have a functional CPC, we searched for Borealin and Survivin homologs in the Arabidopsis genome. Given that the putative INCENP homolog of Arabidopsis, WYR, shares only very weak similarities with its animal counterpart (Kirioukhova et al., 2011), we expected the same to be the case for Borealin and Survivin. Indeed, standard BLAST searches did not result in the identification of likely candidates. Therefore, we made use of the fact that Borealin is transcriptionally controlled by the tumor suppressor protein Retinoblastoma in animals (Cam et al., 2004; Date et al., 2007). Mining a dataset of the genome-wide binding sites of the Arabidopsis Retinoblastoma homolog RETINOBLASTOMA-RELATED (RBR1), revealed several unknown genes likely involved in cell division control (Bouyer et al., 2018) (Sup Fig. 1). One of these genes, *AT4G39630*, showed a weak similarity to Borealin and will be referred to as *BOREALIN-RELATED* (*BORR*) (Supplemental Fig. S1A).

**Figure 1.**
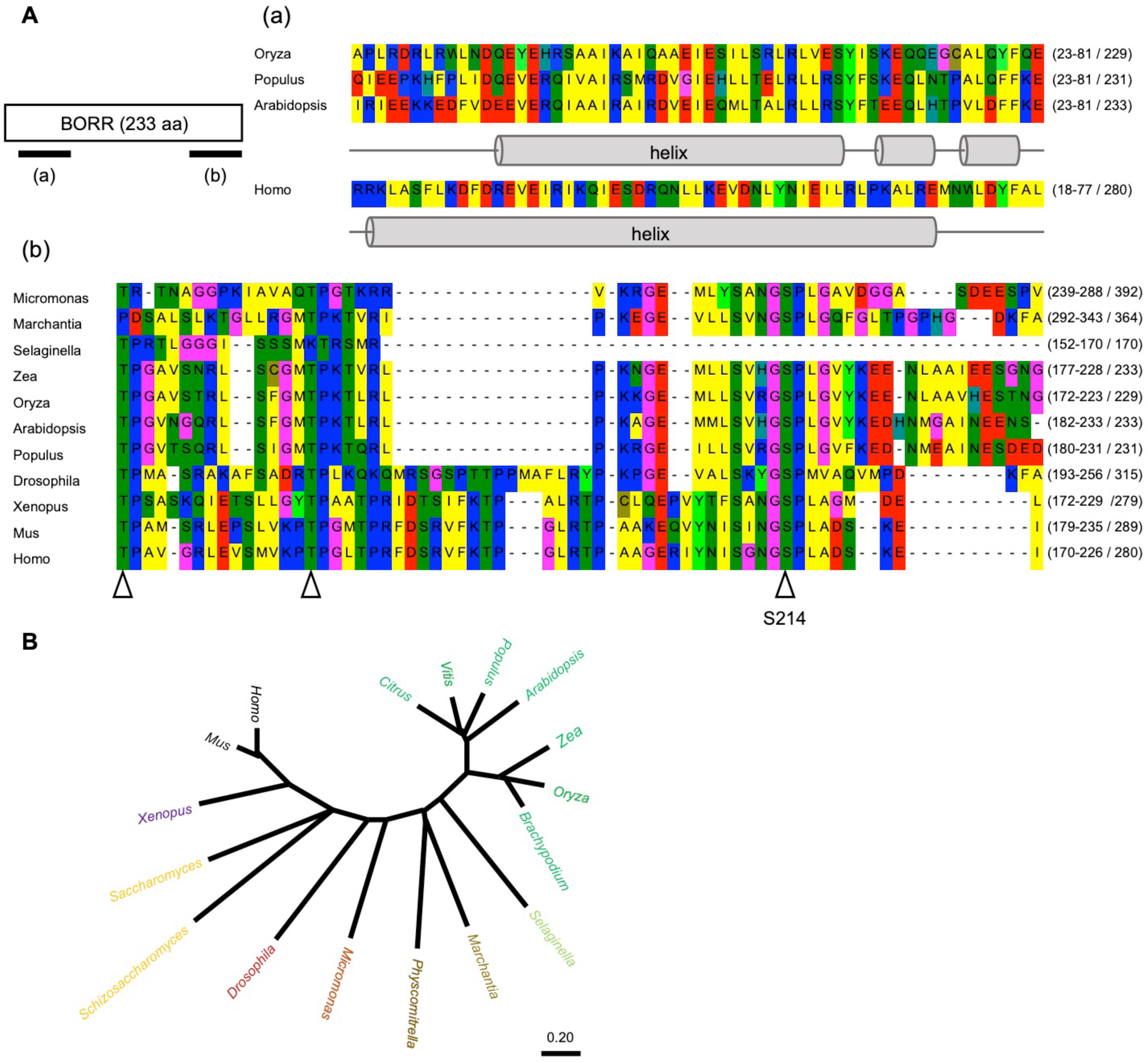
*BOREALIN-RELATED* (BORR) gene structure in plants. A, Protein sequence of BORR in Arabidopsis. Predictions of the N-terminal helix in BORR are shown in (a) and and alignment of the most sequence conserved region of the protein in (b). Arrow heads in (b) indicate the conserved CDK consensus sites. B, Phylogenetic analysis of the Borealin family in yeast, animals and plants. The tree was constructed using MEGA X by the neighbor-joining method.

The C–terminal region of the corresponding protein shows 37% similarity with the central region of human Borealin, which includes highly conserved consensus sites for CDK phosphorylation (Fig. 1A). A second stretch of homology can be found in the N-terminus, which is predicted to adopt an alpha-helical structure. In animals and yeast this N-terminal alpha-helix forms a three-helical bundle with INCENP and Survivin (Fig. 1A) (Jeyaprakash et al., 2007). Notably, the Arabidopsis BORR is considerably shorter than the human homolog (233 aa versus 280 aa), partly as the result of a truncated C-terminal domain. Using *BORR* as a template, we found putative Borealin genes in all branches of the plant kingdom, including angiosperms, gymnosperms, pteridophytes, bryophytes, and algae (Fig. 1B - phylogenetic tree).

### Phenotypic analysis of *borr* mutants

Since no mutants for *BORR* were available in the public mutant collections of Arabidopsis, we generated a mutant by CRISPR/Cas9. The resulting *borr-1* allele has a T insertion in the 2^nd^ exon, i.e. between nucleotides 280 and 281 downstream of the start codon leading to a stop codon and a predicted truncated protein of 67 amino acids (Supplemental Fig. S1A). While the heterozygous *borr-1* plants grow as the wildtype, we could never obtain homozygous mutant plants. Consistently, we observed aborted seeds and undeveloped ovules in siliques of *borr-1 +/-* (Fig. 2A,B).

**Figure 2.**
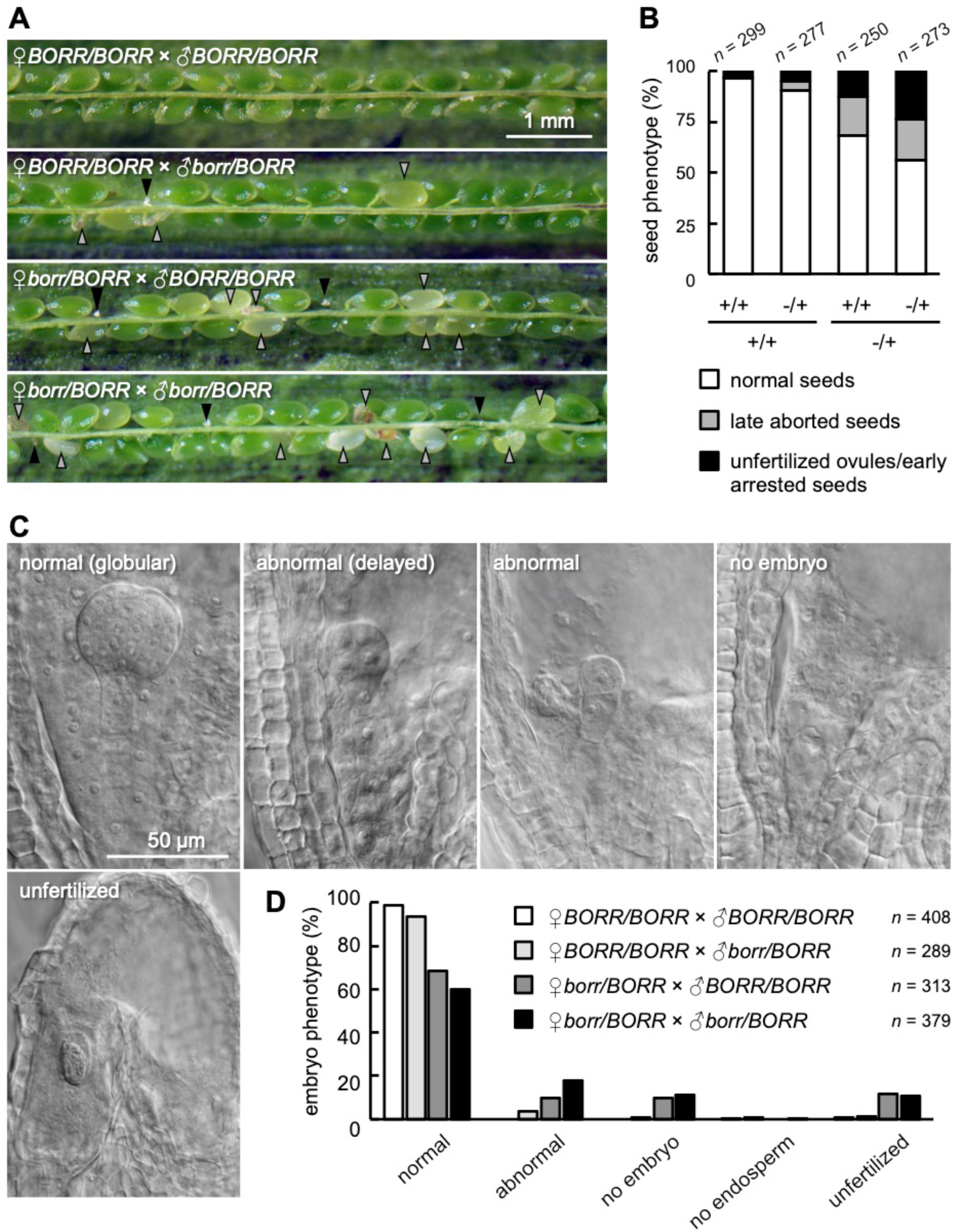
*BORR* is required for seed development. A, Wildtype or heterozygous *borr* mutant plants pollinated with pollen from wildtype or heterozygous *borr* mutants. Black and gray arrowheads indicate tiny white ovules (unfertilized ovules/early arrested seeds) and collapsed brown seeds or seeds without green embryo (late aborted seeds), respectively. B, Proportion of the seed phenotypes shown in (A). Total number (n) of seeds and ovules from five hand-pollinated siliques are shown in each cross. C, Seeds and ovules from the heterozygous *borr* siliques cleared in chloral hydrate solution. At the globular embryo stage, developing seeds with abnormal embryos, developing seeds with no embryo, and unfertilized ovules were observed. D, Classification of the embryo phenotype for each cross. Total number (n) of developing seeds and unfertilized ovules from 6-8 hand-pollinated pistils are shown.

To assess whether this lethal phenotype was indeed due to the frameshift mutation in the *BORR* gene, we constructed reporter lines in which the genomic region of *BORR* was fused to an ORF encoding for GFP. Expression of either an N-terminal (GFP:BORR) or C-terminal (BORR:GFP) GFP fusion to BORR complemented the lethal phenotype of *borr-1* mutant plants corroborating that loss of BORR affects plant reproduction.

To address the nature of the lack of homozygous *borr-1* mutants, we conducted reciprocal crosses of heterozygous mutants with the wildtype. When we used *borr-1* +/- as male parent, the transmission rate was determined to be 44%. Consistently, 96% of *borr1+/-* pollen resembled wild-type pollen and based on DAPI-staining, contained two sperm cells.

In contrast, when we used *borr-1* +/- as female parent, the transmission rate was reduced to 29%, indicating that *BORR* is especially needed for the development and/or function of the female gametophyte. However, when analyzing the mature female gametophyte no obvious developmental defects were observed suggesting that the reduced transmission of *borr-1* through the female gametophyte originated shortly before or after/during fertilization. In accordance with this, we found 12% unfertilized ovules/early arrested seeds and 19% late aborted seeds when we used *borr-1 +/-* as the female (n=250) in contrast to 3% unfertilized ovules/early arrested seeds and 0.3% late aborted seeds in control crosses when we used the wildtype as the female and male parent (n=299)(Fig. 2A,B).

In addition, we observed abnormal embryo development, including delayed and distorted embryos or embryo-like structures in approximately 10% of all ovules/seeds analyzed when *borr-1+/-* was used as the female parent in crosses with the wildtype as a male versus 4% in the control crosses supporting a female gametophytic effect of *borr* (n=313 and n=408 for the control cross) (Fig. 2C,D). One likely explanation for this is that the divisions leading to the development of the embryo sac cause aneuploidy in *borr* that preclude and/or severely interfere with embryo development.

In addition, *BORR* is needed during embryo development since the number of seeds with embryonic defects almost doubled in self-fertilized *borr-1* mutants (Fig. 2D). Thus, BORR is an essential gene needed for cell proliferation and development during early stages of the plant life cycle.

To address the *BORR* function after embryo development, we generated *BORR* knockdown plants by expressing two artificial microRNAs (amiRNA) targeted against the 2^nd^ (*amiBORR#1*) and 3^rd^ exon (*amiBORR#2)* of *BORR*, respectively (Fig. 3A). Most transgenic plants expressing *amiBORR#1* (19 out of 25) and *amiBORR#2* (15 out of 20) showed a dwarf phenotype. For the following analyses, we selected two transgenic plants for each construct (*amiBORR#1-1*, *amiBORR#1-2*, *amiBORR#2-1* and *amiBORR#2-2*). All four knockdown plants had a similar level of *BORR* transcript reduction (Fig. 3B) and had a dwarf phenotype with curled leaves during the vegetative stage (Supplemental Fig. S2A). At the flowering stage, all *BORR* knockdown plants were bushy and exhibited a typical *bonsai* phenotype (sometimes also called *broom stick* phenotype), which is commonly observed in mutants with low APC/C activity and is characterized by short inflorescences, which are often curled at the very end with only a few developing siliques (Supplemental Fig. S2B and C) (Saze and Kakutani, 2007; Zheng et al., 2011).

**Figure 3.**
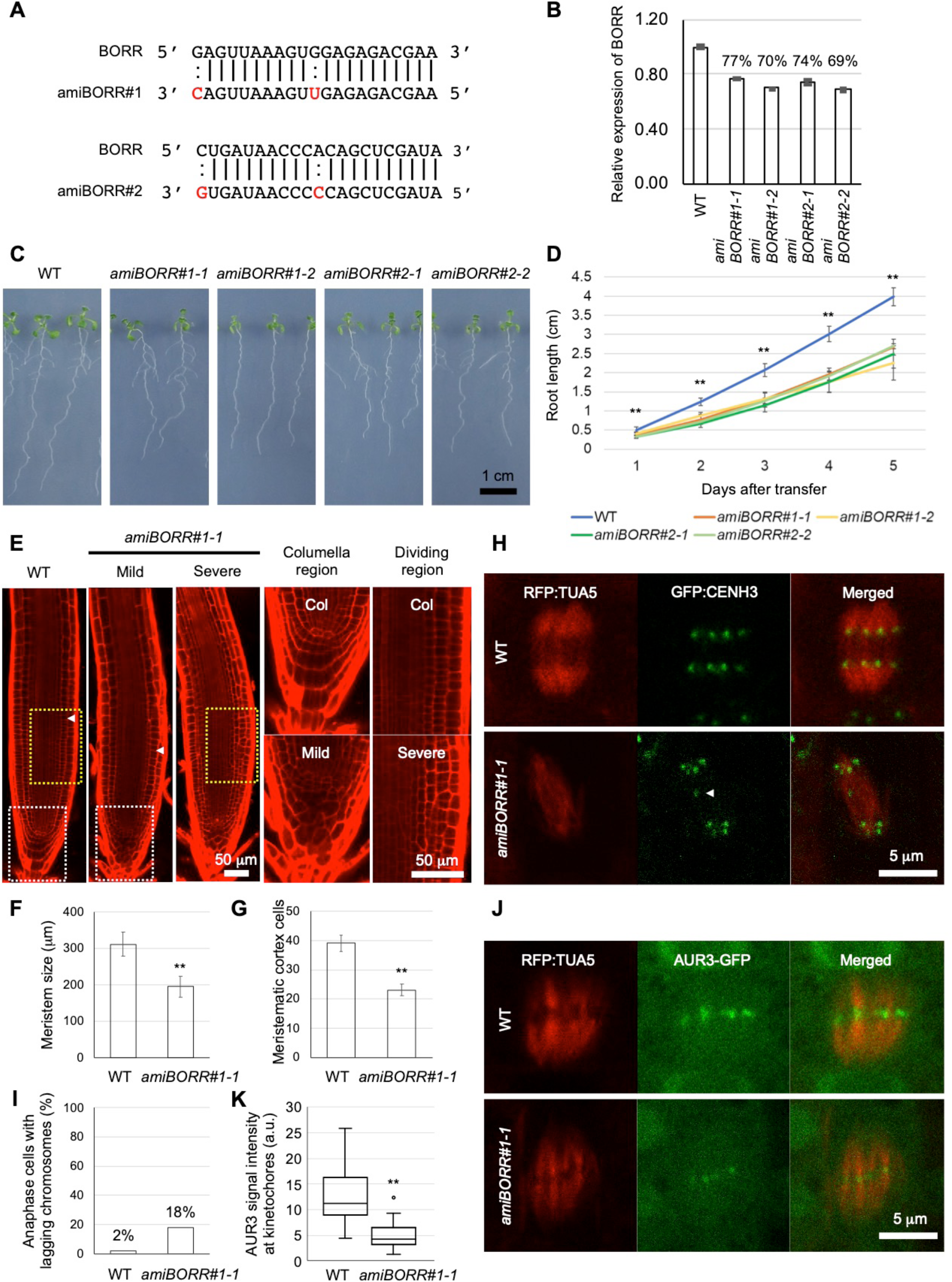
Thenotypical analysis of artificial miRNA-mediated *BORR* knockdown plants. A, Sequence alignments of artificial miRNAs target sites on *BORR* mRNA. B, Relative expression level of *BORR* in the *amiBORR* plants was confirmed by qRT-ysis with three biological replicates. C, 9-day-old wild-type and *amiBORR* seedlings. D, Root growth measurements o ype and *amiBORR* plants. 4-day-old seedlings were transferred to a MS plate and root lengths were measured for 5 day .01, Student’s t-test. Error bars indicate SD *(n* = 10). E, Confocal images of 7-day-old wild-type and *amiBORR#1-1* roots sta propidium iodide. *amiBORR#1-1* roots were categorized into ‘Mild’ and ‘Severe’ groups based on the phenotype. Ar:ate the boundary between the dividing region and the elongation region of the root. The Panels *Columella region* and *Divi >n* show close-up views of the regions marked by white and yellow boxes, respectively. F and G, Root meristem size (F) ber of meristematic cortex cells (G). **P<0.01, Student’s t-test. Error bars indicate SD *(n* = 15). H, Representative imag, ally distributed and lagging chromosomes in 7-day-old WT and *amiBORR#1-1* root cells. Microtubules and centromeres: ilized by TagRFP:TUA5 and GFP:CENH3, respectively. The arrow indicates a lagging chromosome. I, Frequency of lag mosomes in anaphase in **(H).** Error bars indicate SD *(n* = 50). J, Representative images of AUR3 accumulation leve toch ore s in 7-day-old wild-type and *amiBORR#1-1* root cells. Microtubules and AUR3 are visualized by TagRFP:TUA5‘.3: GF P, respectively. K, AUR3 signal intensity at kinetochores in (J). 40 AUR3-GFP signals at kinetochores from 10 cells sured. Median (center line), IQR (box), 1.5 x IQR (error bars), and outliers (circles) are shown. **P<0.01, Student’s t-test. indicate SD.

Primary root growth was also compromised in all *BORR* knockdown plants (Fig. 3C and D). Microscopic analyses of *amiBORR#1-1* revealed that the root meristem size was reduced (Fig. 3E-G). Moreover, an aberrant pattern of cell divisions was found in the columella of all roots analyzed (25 out of 25). In 40% of all seedlings (10 out of 25), altered division patterns were also present in the epidermal, cortex and endodermal layers underlining that *BORR* is required for proper cell division (Fig. 3E).

To assess whether Arabidopsis *BORR* has a function in chromosome segregation as well, *amiBORR#1-1* was introgressed into a transgenic line expressing both a microtubule (RFP:TUA5) and a centromere (GFP:CENH3) marker (Komaki and Schnittger, 2017). Indeed, we could frequently observe lagging chromosomes in *amiBORR#1-1* plant cells (9 out of 50), a phenotype that hardly occurred in the wildtype control plant cells (1 out of 50) (Fig. 3H and I). To examine whether the lagging chromosomes are related to a compromised AUR3 localization, we introduced a previously generated *AUR3:GFP* reporter into *amiBORR#1-1* plants (Komaki and Schnittger, 2017). While some AUR3:GFP signal could be still detected at the kinetochores in *amiBORR#1-1* plants, the signal intensity was much weaker than in wild-type plants, suggesting that Arabidopsis BORR ensures chromosome segregation through AUR3 localization (Fig. 3J and K). Taken together, corresponding to Borealin function in animals, *BORR* is required for proper chromosome segregation and cell division in Arabidopsis.

### Interaction scheme of CPC components in Arabidopsis

To reveal the molecular network of the Arabidopsis CPC, we investigated the interaction of BORR with INCENP and the three Aurora kinases of Arabidopsis. Although Borealin has a conserved coiled-coil domain, which is known as an INCENP binding site in other organisms (Jeyaprakash et al., 2007), an interaction between BORR and INCENP was not detected by a yeast two-hybrid interaction assay (Y2H assay) (Fig. 4A). We next performed an *in vivo* co-immunoprecipitation (IP) assay. To this end, anti-GFP immuno-precipitates from total protein extracts of plants expressing both *BORR:RFP* and *GFP:INCENP* or from plants expressing *BORR:RFP* and free *GFP* as a negative control were probed with an anti-RFP antibody. BORR:RFP was only detected in the extract of plants co-expressing GFP-INCENP, indicating that Arabidopsis BORR could be part of a CPC *in vivo* (Fig. 4C).

**Figure 4.**
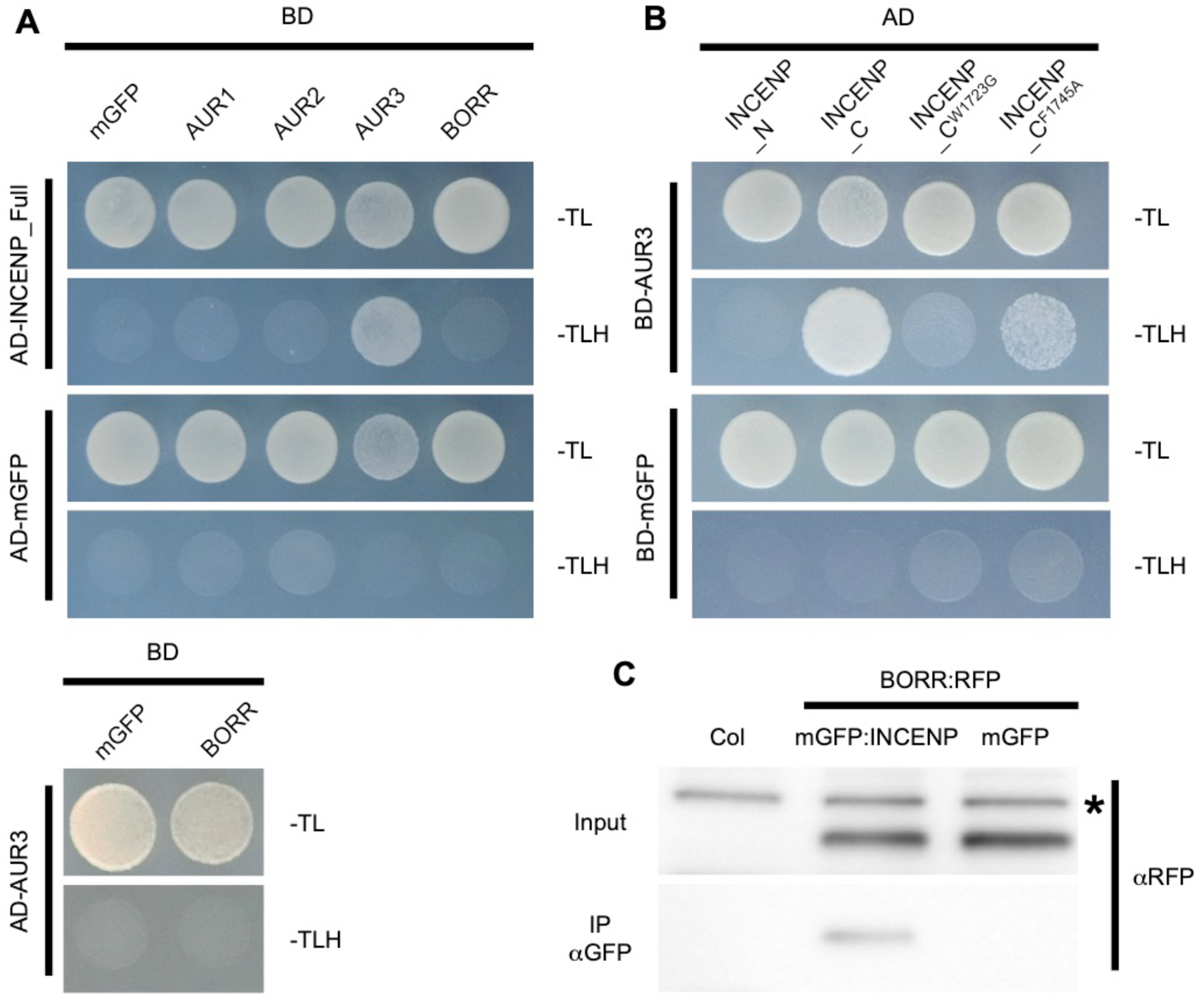
Interaction among the CPC components. A, Interaction among the CPC components were revealed by yeast two-hybrid assays. B, Interaction between AUR3 and various regions of INCENP or IN-box­ mutated INCENP. Monomeric GFP (mGFP) was used as a negative control. Each strain was spotted on SD plates without tryptophan and leucine (-TL; control media) or without tryptophan, leucine, and histidine (-TLH; selection media) and photographed after incubation at 30°C for 2 days. C, Co­ immunoprecipitation of INCENP with BORR. 7-day-old Arabidopsis seedlings expressing BORR:RFP and mGFP:INCENP or BORR:RFP and mGFP were used for IP with an anti-GFP antibody. Both input and IP fraction were subjected to immunoblotting with an anti-RFP antibody. The asterisk on the input panel indicates a nonspecific band.

Consistent with the topology of the CPC in animal and yeast, we further found that INCENP interacts specifically with AUR3 through its C-terminal region while no interaction was detected with AUR1 or AUR2 (Fig. 4A, B). The C-terminal region of INCENP contains an Aurora B binding domain, called IN-box, which is conserved from yeast to mammals (Adams et al., 2000). Since Arabidopsis INCENP also possesses a putative IN-box in its C-terminus, we exchanged Trp1723 to Gly and Phe1745 to Ala. These residues are conserved in yeast and animals, and the

Trp1723 mutation leads to loss of interaction with Aurora B in *in vitro* assays (Sessa et al., 2005), as well as in chicken culture cells (Xu et al., 2009). Consistent with a key role of the putative IN-box, Arabidopsis INCENP^F1745A^ partially and INCENP^W1723G^ completely failed to interact with AUR3 (Fig. 4B).

### Expression pattern and subcellular localization of the CPC components in mitosis

To investigate the spatial and temporal expression pattern of *BORR*, we generated transgenic plants harboring a genomic fragment of the *BORR* gene fused to a beta-glucuronidase (GUS) gene before the STOP codon. As expected, strong GUS activity was observed in both the shoot and root meristems of seedlings, indicating that *BORR* is expressed in proliferating cells (Fig. 5A-C). In addition, *BORR* was also found to be strongly expressed in flowering tissues including both the male and female reproductive organs (Fig. 5D-F).

**Figure 5.**
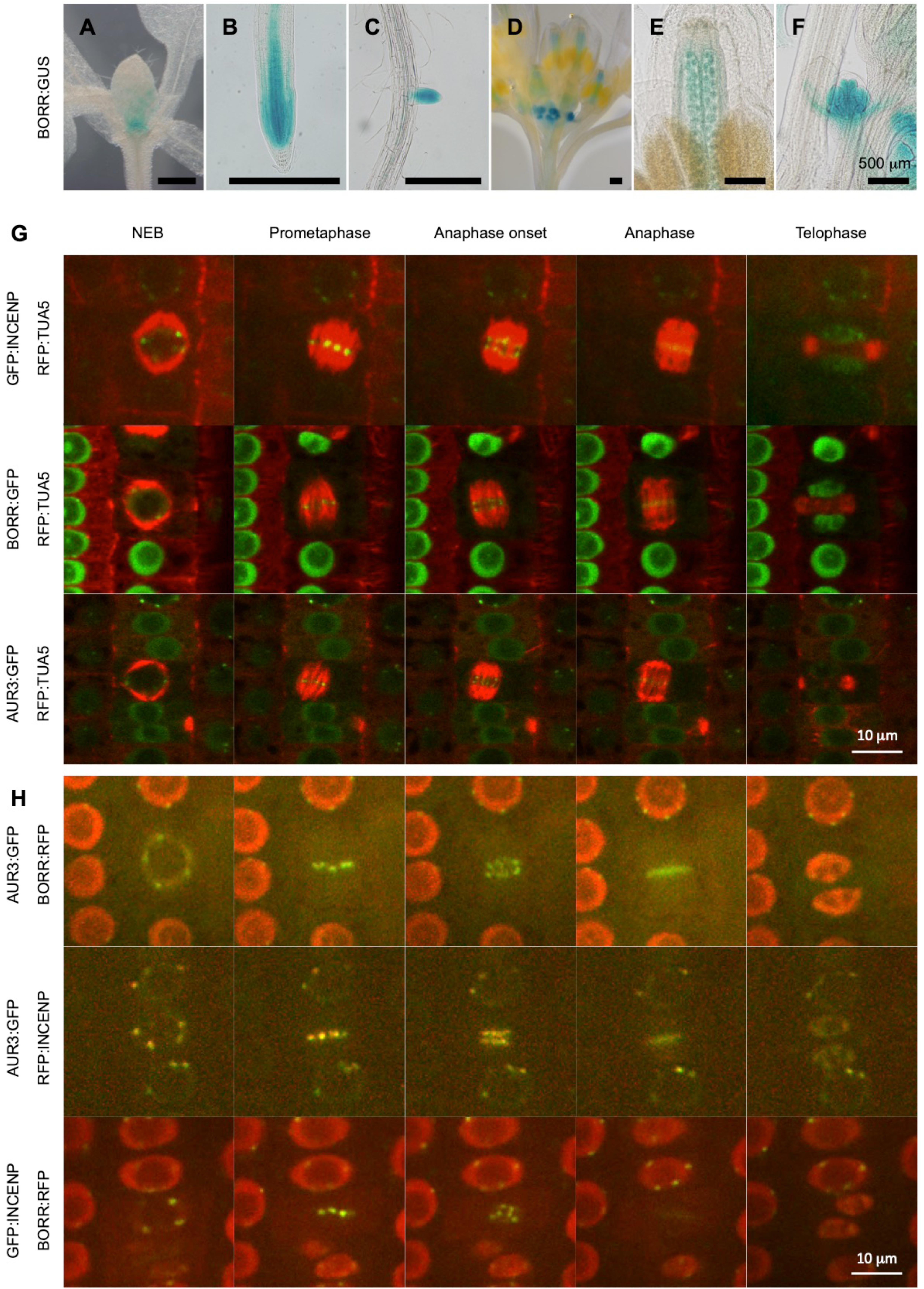
Expression and subcellular localization of CPC components. A to F, Expression pattern of BORR:GUS in the shoot (A), root (B), lateral roots (C), inflorescence (D), ovules (E), and young flower buds (F). 10-day­ old seedlings were used for (A) to (C), and 4-week old plants were used for (D) to (F). Three independent transgenic lines were analyzed and representative images are shown. G, Subcellular localization of GFP:INCENP, BORR:GFP, and AUR3:GFP during the cell cycle. Each reporter line was crossed with TagRFP:TUA5-expressing plants to visualize microtubule structures. H, Co-localization of the CPC components during the cell cycle. For live imaging of (G) and (H), root tips of 5-day-old seedlings were used.

To reveal the subcellular localization of the CPC components in plants, we made use of the functional BORR reporter line used for the complementation studies above. In addition, we constructed reporter lines in which the genomic region of *INCENP* was fused to an ORF encoding for GFP. In contrast to BORR, only the N-terminal (GFP:INCENP) GFP fusion of INCENP could complement the lethal phenotype of *incenp* homozygous mutants (Supplemental Fig. S1D and E). For AUR3, we used a previously published reporter line (*AUR3:GFP*) in the wildtype background (Komaki and Schnittger, 2017).

First, we crossed the three CPC reporter lines with *RFP:TUA5* expressing plants to check the localization of CPC components during mitosis. All three CPC components showed the same localization pattern (Fig. 5G; Supplemental Movies S1-3). In interphase, they localized to the nucleus. Before nuclear envelope breakdown (NEB), the CPC components strongly accumulated at the kinetochores until anaphase onset. Once chromosomes moved toward the spindle poles, they localized to the middle part of the phragmoplast. Interestingly, in early telophase, the CPC components moved back to the nucleus even though there still was an expanding phragmoplast.

To corroborate that all three CPC components co-localize, we created transgenic plants which expressed BORR:RFP and GFP:INCENP, BORR:RFP together with AUR3:GFP and RFP:INCENP along with AUR3:GFP (Table 1). Microscopical analyses revealed a tight co-localization pattern through the cell cycle in all three lines, suggesting that BORR, INCENP and AUR3 work together as a plant CPC (Fig. 5H; Supplemental Fig. S3 and 4; Supplemental Movies S4-6).

**Table 1.**
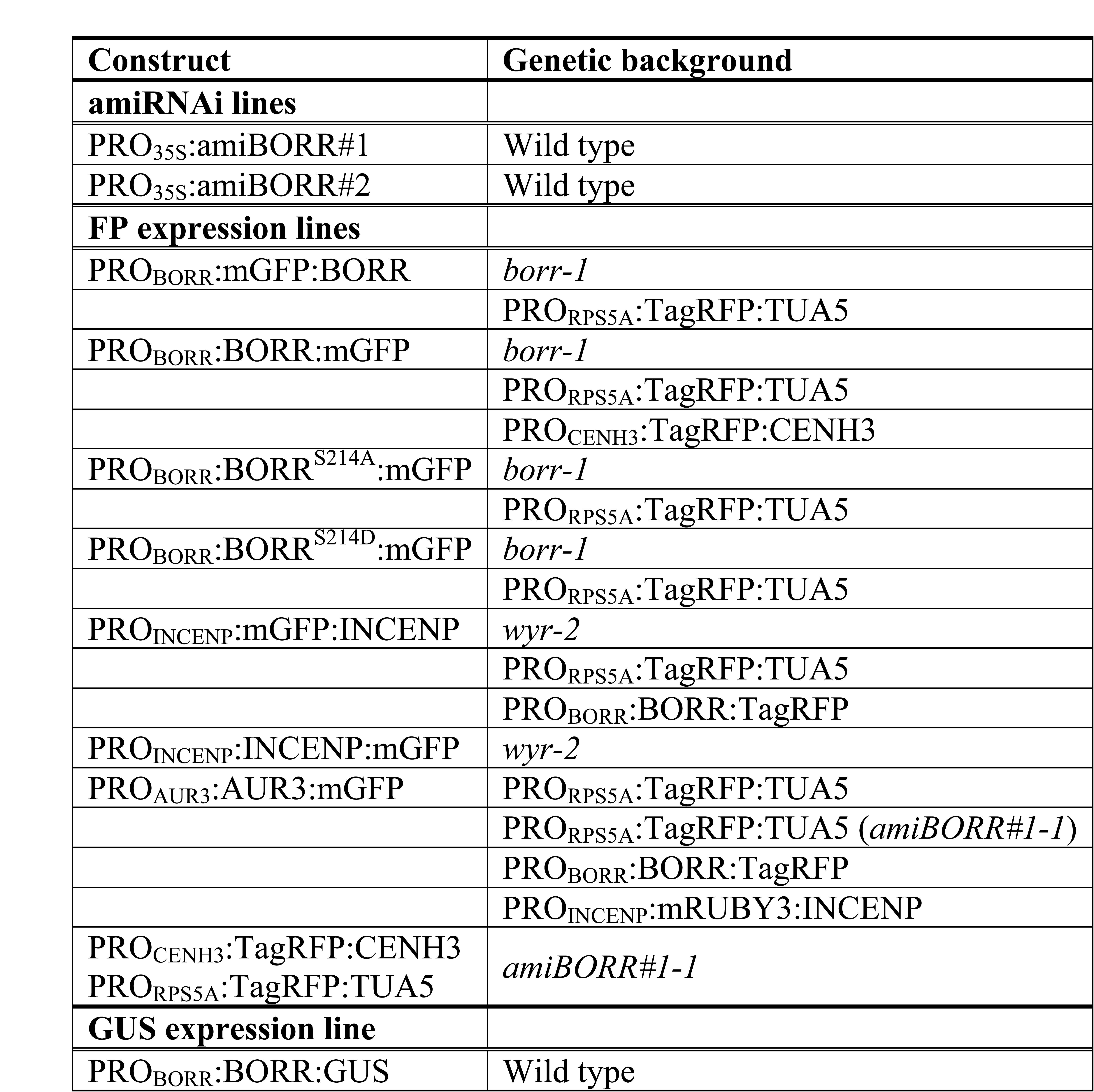
The genetic material generated in this study.

In other organisms, it has been reported that the CPC localizes to the inner centromere region to monitor kinetochore-microtubule attachments (Hindriksen et al., 2017). To reveal the localization of the CPC precisely, we crossed the *BORR:GFP* line with the inner kinetochore marker *RFP:CENH3*. Just after NEB, these two fluorescent reporters showed a high level of overlapping signal. After prometaphase, BORR:GFP localized closer to the inner region of the kinetochore than RFP:CENH3. Yet, both proteins still overlapped in their localization pattern. Shortly before anaphase onset, RFP:CENH3 formed two lines along the metaphase plate, and BORR:GFP localized between these two lines with no overlap (Fig. 6; Supplemental Movie S7). This result demonstrates that the CPC localizes to the inner centromere region in plants.

**Figure 6.**
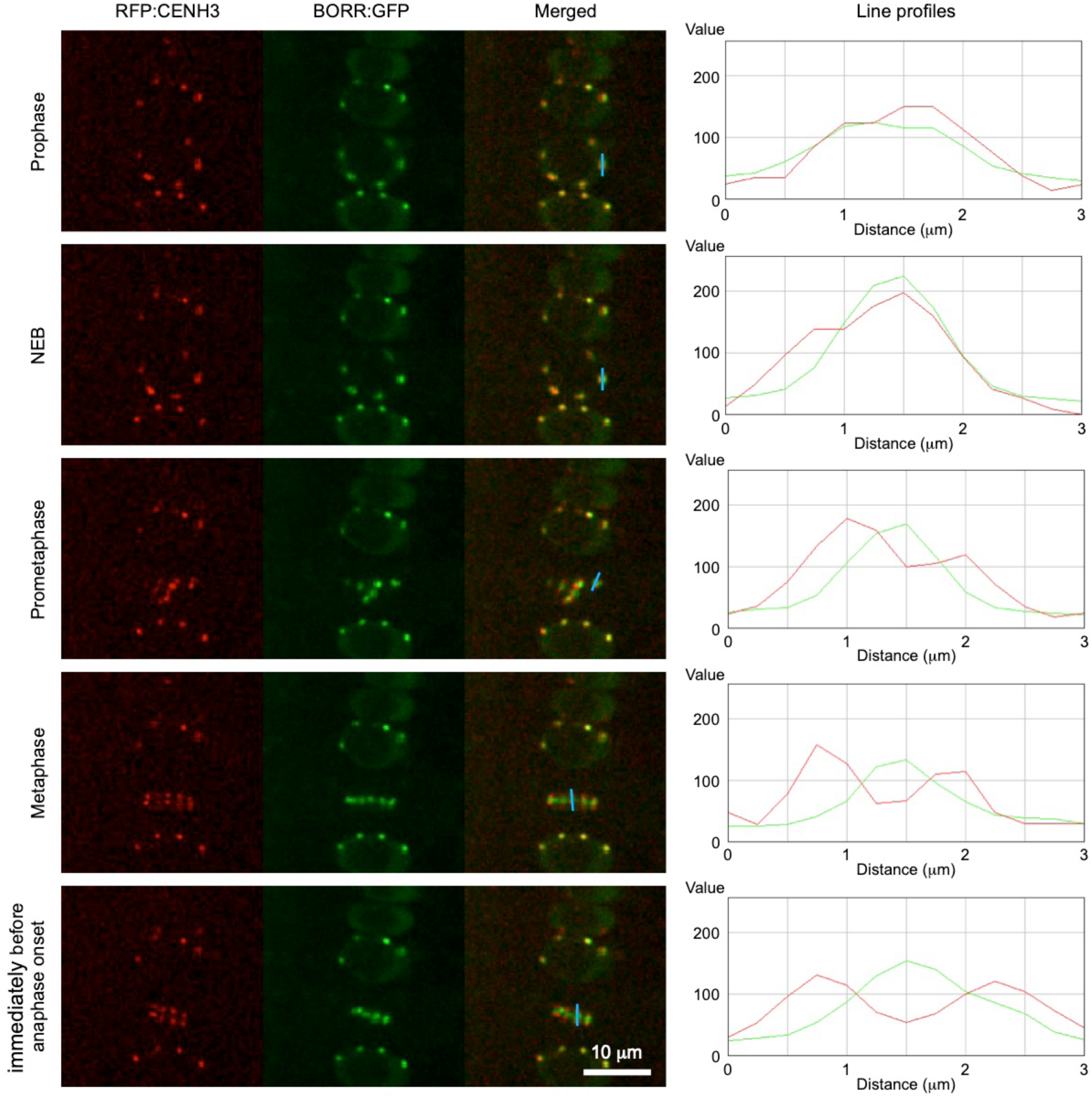
CPC localizes to inner centromeres. Localization of the CPC from prophase to immediately before anaphase onset. Inner kinetochores and CPC complexes are visualized by TagRFP:CENH3 and BORR:GFP, respectively. For live cell imaging, root tips of 5-day-old seedlings were used. Blue bar indicates the position where the line profiles were obtained.

In humans, Borealin function is regulated by phosphorylation. Especially the phosphorylation of S219, a putative Cdk1 target residue, in the central region of the protein affects its stability and centromere localization (Kaur et al., 2010; Date et al., 2012). Since this Cdk1 phosphorylation site resides in the most conserved region between human and plant Borealin is conserved in BORR (Figure 1B), we generated a phospho-mimic (BORR^S214D^:GFP) and a dephospho variant (BORR^S214A^:GFP) (Dissmeyer and Schnittger, 2011), and transformed them into the heterozygous *borr-1* mutants to evaluate their functionality. Notably, both constructs could complement the lethal phenotype of *borr-1* mutant plants, and showed a normal BORR localization pattern (Supplemental Fig. S1A-C). Thus, the physiological importance of this conserved CDK target motif has yet to be resolved in plants.

### Expression pattern and subcellular localization of the CPC components in meiosis

Since the plant meiotic spindle checkpoint seems to be less stringent (Wijnker and Schnittger, 2013; Komaki and Schnittger, 2016), we asked whether BORR and INCENP are present in meiosis. Analyzing male meiosis, we found that both proteins localized to kinetochores until the onset of anaphase I (Fig. 7A and B; Supplemental Movies S8 and S9). Interestingly, although no cytokinesis occurs after meiosis I in Arabidopsis meiocytes, both proteins localized to the division plane shortly after anaphase I either representing a late anaphase midzone or a phragmoplast midzone-like structure (Fig. 7A and B; Supplemental Movies S8 and S9). The transition between both phases is currently only poorly defined but the localization of BORR and INCENP could possibly contribute to the understanding of the composition and dynamics of these structures.

**Figure 7.**
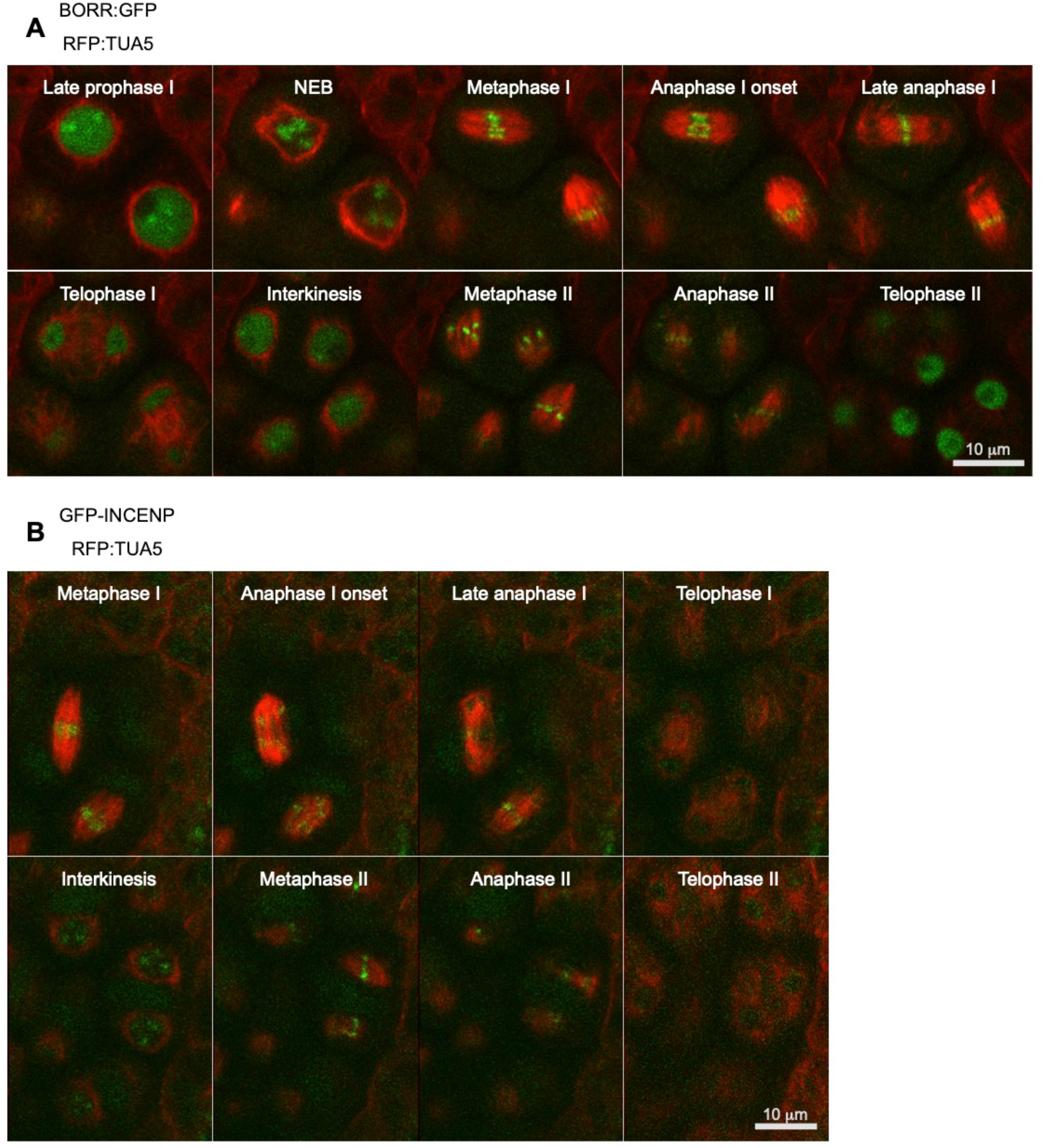
Subcellular localization of CPC components in meiosis. A and B, Subcellular localization of BORR:GFP (A) and GFP:INCENP (B) during meiosis. Each reporter line was crossed with TagRFP:TUA5-expressing plants to visualize the microtubule structures. For live imaging of (A) and (B), flower buds of 1-month-old plants were used.

The localization of BORR and INCENP in the second meiotic division resembled the localization of both proteins in mitosis, i.e. at the kinetochores and subsequently at the phragmoplast (Fig. 7A and B; Supplemental Movies S8 and S9).

## Discussion

The CPC is essential for proper cell division in animals and yeast. Whether CPC activity exists in plants and how the potential complex is composed was up to the presented work not clear. Here, we have identified and characterized *BORR*, a Borealin homolog in Arabidopsis. While the presence of a functional homolog of *Survivin* in plants is currently still unclear, the existence of *BORR* together with localization data for AUR3 and an INCENP homolog demonstrates that a CPC is present in plants and of equal importance as in animals.

BORR and INCENP localize to kinetochores in mitosis and meiosis. Both proteins could also be co-precipitated from seedlings indicating that BORR and INCENP indeed work in one complex. Consistent with previous reports, we observed that AUR3 accumulates at mitotic kinetochores (Fig. 5G; Supplemental Movie S3) (Demidov et al., 2005), while AUR1 and AUR2 localize to the mitotic spindle (Van Damme et al., 2011). In addition, out of the three AUR proteins in Arabidopsis, only AUR3 interacted with INCENP in our Y2H assays (Fig. 4). These data suggest that only AUR3 acts as the catalytic subunit of the CPC at the inner centromere in plants.

In animals, the CPC localizes to inner centromere to monitor inter-kinetochore tension. Since proper kinetochore-microtubule attachments are not established during prophase, the distance between inner centromere and kinetochores is very small allowing the CPC-dependent centromere-localized Aurora B activity to act on kinetochores and by to destabilize erroneous attachments of microtubules.

Once proper attachments are formed and inter-kinetochore tension is built up, Aurora B localized to inner centromere is spatially separated from kinetochores leading to the formation of a stable bipolar spindle. Since we found that the kinetochore signals moved away from BORR signals during cell-cycle progression, we propose that the plant CPC also acts as a tension sensor.

At anaphase onset, the animal CPC translocates from kinetochores to the cell division plane. Although plant cell division is strikingly different (Mü ller and Jü rgens, 2016), we could show that Arabidopsis BORR, INCENP and AUR3 also accumulate at the division plane at the beginning of cell division (Fig. 5G; Supplemental Movie S1-3). Interestingly, they accumulate the reforming nuclei at telophase before the expanding phragmoplast is completely disassembled, i.e. cytokinesis is finished, while in animals the CPC stays at the cleavage site until the two daughter cells are formed (Carmena et al., 2012).

Remarkably, previous studies revealed that the two alpha-Aurora members, AUR1 and AUR2, localize to the division plane until the end of cytokinesis (Demidov et al., 2005; Kawabe et al., 2005). Therefore, the plant CPC might be needed for the initiation but may not be necessary for later steps of cell division. In animals, TPX2 recruits Aurora A to the division plane (Kufer et al., 2002). However, TPX2 does not localize to the division plane in plants although it and its homologs also interact with alpha-Aurora members in Arabidopsis (Petrovská et al., 2012; Boruc et al., 2019). Thus, it is still not clar how alpha-Aurora localization to the division plane is controlled.

It seems likely that there is a yet unidentified interaction partner of the CPC in plants that has affinity to the plus-end of microtubules and causes the localization of the CPC, i.e. AUR3 to kinetochores. After separation of the chromosomes in anaphase, this or a different factor then promotes the accumulation in the spindle midzone/early phragmoplast.

Since each component of the CPC is needed for its activity, loss of any CPC component leads to the same mutant phenotypes as seen in Aurora B mutants (Honda et al., 2003; Vader et al., 2006). CPC mutants typically exhibit cell division defects and lagging chromosomes resulting from incorrect microtubule-kinetochore attachments. These phenotypes are frequently coupled with ploidy changes causing cancer in mammals (Tang et al., 2017). Notably, a complete loss-of-function of any of the CPC components leads to lethality (Cutts et al., 1999; Uren et al., 2000; Lu et al., 2008; Yamanaka et al., 2008). The sporophyte of *borr-1* heterozygous knockout plants grew like the wildtype. However, no homozygous *borr* mutants could be recovered and the heterozygous mutants harbor undeveloped ovules, aborted seeds, and embryonic defects in the forming siliques. Similar effects were observed in the *INCENP* mutant *wyr* (Kirioukhova et al., 2011). However, in contrast to *borr*, the loss of INCENP causes an arrest in development of both the female and the male gametophytes. We can currently not rule out whether INCENP has a specific and BORR-independent function during the gametophyte life phase. It also possible that INCENP has a shorter half-life than BORR, given that it is a large protein, and that hence the gametophytes run out of sporophytically inherited protein levels much earlier than it is the case for the much smaller BORR protein.

Using knock-down plants, we could further reveal that reduction of *BORR* results in mitotic defects including lagging chromosomes and abnormal cell division which appear to be likely caused by a compromised localization of AUR3. A previous study reported that hesperadin treatment, which inhibits the AUR3 kinase activity *in vitro*, induces lagging chromosomes in tobacco BY-2 cells (Kurihara et al., 2006), suggesting that the AUR3 function in chromosome segregation is conserved in the plant lineage.

Interestingly, the *BORR* knockdown plants showed a typical *bonsai* phenotype, which is characterized by the inhibition of internode elongation and the premature termination of the shoot apical meristem. Although the molecular mechanism is still not known, the *bonsai* phenotype is associated with a reduction of APC/C activity (Saze and Kakutani, 2007; Zheng et al., 2011).

The SAC is another M-phase checkpoint, which works together with the CPC to ensure faithful chromosome segregation. The primary role of the SAC is delaying APC/C activity until all kinetochores are properly attached to the spindle microtubules. Therefore, one possible explanation of the *bonsai* phenotype is that the SAC is over-activated in *BORR* knockdown plants. Indeed, these two M-phase checkpoints are directly or indirectly connected in other organisms (Trivedi and Stukenberg, 2016). Further studies are needed to understand the relationship between SAC and CPC in plants.

The final non-catalytic CPC subunit, Survivin, remains elusive in plants. Survivin localizes to the inner centromere upon phosphorylation of histone H3 at Thr3, which in animals is catalyzed by the Haspin kinase (Kelly et al., 2010). Survivin localization is required for recruiting the entire CPC to the inner centromere. Therefore, inhibition of Haspin kinase activity leads to the dissociation of the CPC from inner centromere in mammals. Recently, it was shown that inhibition of Haspin kinase activity with 5-iodotubercidin induces the disruption of AUR3 localization at the inner centromere in BY-2 tobacco culture cells as well (Kozgunova et al., 2016). Although this result indicates that Haspin kinase activity is important for proper AUR3 localization at the inner centromere, no Survivin homolog could be identified in plants on a sequence level. However, a functional homolog might still exist. Alternatively, plants might employ a different mechanism for CPC localization.

The phosphorylation status of its different components is of key importance for the regulation of the CPC in animals and yeast. For instance, yeast INCENP is cooperatively phosphorylated by Cdk1 and Aurora, which prevents the CPC binding to the spindle before anaphase (Goto et al., 2006; Nakajima et al., 2011). In addition, Casein kinase 2 phosphorylates the human Survivin, which leads to its exclusion from the nucleus in interphase (Barrett et al., 2011). Borealin is also phosphorylated by many kinases, including Cdk1 that is required for the targeting of the CPC to kinetochores (Kaur et al., 2010; Date et al., 2012). However, the mutation of a conserved CDK phosphorylation site neither obviously altered BORR localization nor reduced the activity of the protein in a way that it would result in a mutant phenotype. Hence, further work is needed to shed light on the regulation of the plant CPC with the present study opening the door for an in-depth analysis of the CPC function in plants with focus on kingdom-specific aspects of its regulation and activity.

## MATERIALS AND METHODS

### Plant Materials and Growth Conditions

The *Arabidopsis thaliana* accession Columbia (Col-0) was used as the wildtype in this study. All mutants are in the Col-0 background. Plants were grown on a solid medium containing half-strength Arabidopsis nutrient solution (Haughn and Somerville, 1986), 1% (w/v) sucrose and 1.5% (w/v) agar in a growth chamber (16h of light; 21°C/8h of dark; 18°C). The T-DNA insertion line GABI_65B09 (*wyr-2*) was obtained from the Nottingham Arabidopsis Stock Center. The *borr-1* line was generated by CRISPR/CAS9 (Fauser et al., 2014). Primer pairs for genotyping are described in Supplemental Table S1 and Supplemental Fig. S1.

### Plasmid Construction and Transgenic Plants

The plasmid construction for the CRISPR/CAS9 system was performed as described in Fauser et al. (2014). To obtain the *borr* knockout plants, 20-bp gene-specific spacer sequences of *BORR* gene (Supplemental Table S1) were cloned into the pEn-Chimera, followed by LR recombination reactions with the destination vector pDe-CAS9. The plasmid construction for the artificial microRNA system was performed as described in Carbonell et al. (Carbonell et al., 2014). To obtain the *BORR* knockdown plants, 75-bp gene-specific sequences of *BORR* gene with *AtMIR390a* backbone (Carbonell et al., 2014) (Supplemental Table S2) were synthesized and cloned into *pDONR221*, followed by LR recombination reactions with the destination vector *pGWB602*. To create the *PRO_BORR_:BORR:GUS* construct, 2 kb upstream of the start codon and 1 kb downstream of the stop codon of the *BORR* gene were amplified by PCR and cloned into *pENTR2B* by SLiCE. The *Sma*I site was inserted in front of the stop codon of the *BORR* construct. The resulting construct was linearized by *Sma*I digestion and was ligated to the *GUS* gene, followed by LR recombination reactions with the destination vector *pGWB501* (Nakagawa et al., 2007). To create *PRO_BORR_:BORR:FPs* constructs, the *GUS* gene in *PRO_BORR_:BORR:GUS* construct was replaced by the ORF for monomeric *GFP* (*mGF*P) or *TagRFP-T*. To create the *PRO_BORR_:GFP:BORR* construct, the *Sma*I site was inserted in front of the start codon of *BORR* construct. To create *PRO_INCENP_:FPs:INCENP*, the genomic fragment of *INCENP* gene was amplified by PCR and cloned into *pENTR2B* by SLiCE method. The *Sma*I site was inserted in front of the start codon of INCENP. The resulting construct was linearized by *Sma*I digestion and was ligated to the *mGFP* or *mRUBY3* gene, followed by LR recombination reactions with the destination vector *pGWB501*. Primer pairs for plasmid construction are described in Supplemental Table S1. Transgenic plants were generated by the floral dip method. The *Agrobacterium tumefaciens* strain *GV3101* (*pMP90*) harboring the gene of interest on a binary plasmid was grown in 3 ml of LB media at 28°C. Agrobacteria were resuspended in 3 ml of transformation buffer containing 5% sucrose and 0.05% silwet L-77, and used for plant transformation.

### Expression Analysis by qRT-PCR

Total RNA was isolated from 7 day-old seedlings with the RNeasy Plant Mini Kit (Qiagen). 300 mg of total RNA were reverse transcribed with ReverTra Ace qPCR RT Master Mix with gDNA Remover (TOYOBO) according to the manufacturer’s instructions. Real-time PCR was performed using the Roche LightCycler 480 and the TB Green Premix Ex Taq (TaKaRa). PP2AA3 (*AT1G13320*) was used as the reference gene (Czechowski et al., 2005). Primer pairs for *BORR* and *PP2AA3* are described in Supplemental Table S1. All experiments were performed in three biological replicates.

### Confocal Microscopy and Image Analysis

Root tips of 5 day-old seedlings were used for live cell imaging. Samples were put on glass bottom dishes and covered with a solid medium containing half-strength Arabidopsis nutrient solution, 1% sucrose and 1.5% agar. Confocal images of mitotic cells were acquired by an inverted Nikon ECLIPSE Ti-U microscope equipped with a YOKOGAWA CSU-X spinning disc detector unit connected to an EM-CCD camera (iXon3 DU897; Andor) and a laser combiner system (500 series; Andor), using a Plan Apo 60x/1.20 water immersion objective. GFP was excited at 488 nm with a 520/35 emission filter and TagRFP-T and mRUBY3 at 561 nm with a 617/73 emission filter. Images were obtained at 20 second intervals and corrected for sample drift using the StackReg plugin (ImageJ version 1.49). To obtain line profile data, images were analyzed by the RGB Profiler plugin (ImageJ version 1.49).

For meiotic live cell imaging, flower buds from 1 month-old plants were used. Sample preparation was performed as described in Prusicki et al., (2019) (Prusicki et al., 2018).

Images were obtained every 1 min for BORR:GFP and every 3 min for GFP:INCENP. Images were corrected for the sample drift by the StackReg plugin (ImageJ version 1.49).

### GUS Histochemical Analysis

Samples were fixed in 90% acetone for 15 min and were washed in 50 mM sodium phosphate buffer. The fixed samples were incubated in GUS solution (50 mM sodium phosphate buffer, pH 7.0, 0.5% (v/v) Triton X-100, 0.5 mM K_3_Fe(CN)_6_, 0.5 mM K_4_Fe(CN)_6_ and 0.5 mg ml^-1^ X-gluc) for 1 h at 37°C. After staining, the samples were cleared in chloral hydrate solution (8g chloral hydrate, 1 mL 100% glycerol, 2 mL distilled water).

### Yeast Two-Hybrid Assay (Y2H)

Y2H assays were performed as described in Komaki and Schnittger (2017). All of the cDNAs tested were amplified by PCR using gene specific primers from cDNA made from total RNA of wildtype Arabidopsis, followed by PCR with universal *attB* primers, and cloned into *pDONR221*. The subcloned *BORR* cDNA was recombined into *pGBT9-C* (DNA-BD), which *GAL4-BD* is fused with C-terminus of *BORR* by LR recombination reactions. The other subcloned cDNAs were recombined into the conventional vector *pGBT9* (DNA-BD) or *pGAD424 (*AD). Primer pairs for plasmid construction are described in Supplemental Table S1.

### Protein Extraction and Co-Immunoprecipitation Assay

1 g of 7 day-old seedlings expressing *BORR:RFP* with *GFP:INCENP* or *GFP* was ground to a fine powder in liquid nitrogen with mortar. Total protein was extracted in 2 mL of extraction buffer containing 50 mM Tris-HCl, pH8.0, 150 mM NaCl, 1% IGEPAL CA-630, and a EDTA-Free Protease Inhibitor Cocktail (Roche) for 30 min on ice, and centrifuged for 10 min at 20,000 g at 4°C. The supernatant was incubated for 1 h on ice with anti-GFP magnetic beads (Miltenyi Biotec), and beads were washed 4 times with extraction buffer on the magnetic field. Then, the beads were boiled in SDS sample buffer to release the proteins. Protein samples were detected with 1:2,000 diluted anti-GFP (A6455; Thermo Scientific) and 1:1,000 diluted anti-RFP (AB233; evrogen) as primary antibodies and subsequently with 1:10,000 diluted anti-rabbit IgG, HRP-linked antibody (NA934; GE Healthcare) as secondary antibody.

### Accession Numbers

Sequence data from this article can be found in Arabidopsis Information Resource (TAIR) – database under the following accession numbers: *AUR1* (*AT4G32830*), *AUR2* (*AT2G25880*), *AUR3* (*AT2G45490*), *BORR* (*AT4G39630*), *INCENP/WYRD* (*AT5G55820*). Borealin protein sequence data of other organisms from this article can be found in the NCBI data libraries under the following accession numbers: *Brachypodium distachyon* (XP_003570304.1), *Citrus clementina* (XP_006428073.1), *Drosophila melanogaster* (NP_609279.1), *Homo sapiens* (NP_001243804.1), *Marchantia polymorpha* (P788.1), *Micromonas commoda* (XP_002503544.1), *Mus musculus* (NP_080836.3), *Oryza sativa* (XP_015650359.1), *Physcomitrella patens* (XP_024357151.1), *Populus trichocarpa* (XP_024458072.1), *Saccharomyces cerevisiae* (AJV31725.1), *Schizosaccharomyces pombe* (CAA22184.2), *Selaginella moellendorffii* (XP_024537708.1), *Vitis vinifera* (XP_002280684.1), *Xenopus tropicalis* (NP_001002902.1), *Zea mays* (NP_001142076.1).

## SUPPLEMENTAL DATA

The following supplemental materials are available.

Supplemental Figure S1. Complementation test of BORR and INCENP mutants.

Supplemental Figure S2. Both BORR and AUR3 localize to inner centromeres.

Supplemental Table S1. Primers used in this study.

Supplemental Table S2. Synthesized oligonucleotides used in this study.

Supplemental Movie S1. Subcellular localization of GFP:INCENP during mitosis.

Supplemental Movie S2. Subcellular localization of BORR:GFP during mitosis.

Supplemental Movie S3. Subcellular localization of AUR3:GFP during mitosis.

Supplemental Movie S4. Co-localization of AUR3:GFP and BORR:RFP.

Supplemental Movie S5. Co-localization of AUR3:GFP and RFP:INCENP.

Supplemental Movie S6. Co-localization of GFP:INCENP and BORR:RFP.

Supplemental Movie S7. Comparison of subcellular localization between BORR:GFP and RFP:CENH3.

Supplemental Movie S8. Subcellular localization of BORR:GFP during meiosis.

Supplemental Movie S9. Subcellular localization of GFP:INCENP during meiosis.

## Acknowledgments

We thank Konstantinos Lampou for critical reading and helpful comments to the manuscript. We are grateful to the University of Hamburg for core funding.

## Footnotes

This work was supported through a DFG grant (SCHN 736/8-1) to A.S., and a JSPS KAKENHI Grant (JP18K45678) to S.K. In addition support of MEXT KAKENHI, Grant-in-Aid for Scientific Research on Innovative Areas (JP19H04864) to S.K. and core funding of the University of Hamburg to A.S. is greatly acknowledged.

